# Dysregulated cell signalling and reduced satellite cell potential in ageing muscle

**DOI:** 10.1101/761023

**Authors:** Ketan Patel, Biggy Simbi, Olli Ritvos, Sakthivel Vaiyapuri, Gurtej K Dhoot

## Abstract

Aberrant activation of signalling pathways has been postulated to promote age related changes in skeletal muscle. Cell signalling activation requires not only the expression of ligands and receptors but also an appropriate environment that facilitates their interaction. Here we first examined the expression of SULF1/SULF2 and members of RTK and the Wnt family in skeletal muscle of normal and a mouse model of accelerated ageing. We show that SULF1/SULF2 and these signalling components, a feature of early muscle development are barely detectable in early postnatal muscle. Real time qPCR and immunocytochemical analysis showed gradual but progressive up-regulation of SULF1/SULF2 and RTK/Wnt proteins not only in the activated satellite cells but also on muscle fibres that gradually increased with age. Satellite cells on isolated muscle fibres showed spontaneous *in vivo* satellite cell activation and progressive reduction in proliferative potential and responsiveness to HGF and dysregulated myogenic differentiation with age. Finally, we show that SULF1/SULF2 and RTK/Wnt signalling components are expressed in progeric mouse muscles at earlier stage but their expression is attenuated by an intervention that promotes muscle repair and growth.

## INTRODUCTION

Often ligands are sequestered in the extracellular matrix (ECM) that need to be liberated for them to activate their receptors. It has previously been shown that de-sulfation of the ECM mediated by SULF1/SULF2 enzymes regulates Wnt signalling during development and regeneration (Ai et al., 2003; Dhoot et al., 2001). Skeletal muscle unlike cardiac muscle can successfully repair itself to restore function due to its endowment with resident stem cells called satellite cells that can be activated upon injury. Satellite cells exist normally in a quiescent state and are located under the basal lamina. However, during periods of muscle growth or following muscle injury they become activated and proliferate to repair the damaged muscle. At this stage the daughter cells either follow a differentiation pathway that leads them to eventually fuse with each other or with an existing fibre (Bischoff, 1986). Alternatively, they revert to a quiescent stem cell state. A satellite cell sub-population is maintained for life through regulated and limited self-renewal but the number of satellite cells decreases with age and those that are present display functional deterioration (Gibson MC and E., 1983; Shefer G1 et al., 2006). A similar situation occurs in numerous chronic diseases of the muscle including Duchenne Muscular Dystrophy where repeated rounds of degeneration and regeneration exhaust the satellite cell population leading to fibrosis.

Regenerative ability is also impaired in ageing muscle. Age-related muscle deterioration manifested by loss of muscle mass and strength is well recognised but the basis of such weakness and possible amelioration is not known. This may result from the reduced number of satellite cells or their declining ability to proliferate or differentiate fully due to changed microenvironment with time. Some studies have shown the total number of satellite cell number to decline with age (Gibson MC et al, 1983; Shefer G et al, 2006, Brack AS et al, 2013) that could limit the regenerative potential of aged muscle but the nature of other contributory factors affecting the loss of muscle mass or reduced muscle regenerative capacity with age is less clear.

Reduced number of satellite cells and the finite regenerative potential of resident satellite cells could nevertheless be exhausted following repeated cycles of injury and regeneration. Dysregulated cell signalling imposed by changes in the satellite cell niche could further impair the muscle regenerative capacity. The regulated activity of such cells on the other hand could preserve satellite cell function for longer. Regulated cell signalling ensures well-orchestrated skeletal muscle growth during fetal and postnatal growth and timely onset of muscle differentiation but re-activation of similar processes is not so well maintained in later life.

The muscle satellite cells enter quiescence upon muscle growth completion accompanied by marked down-regulation of most cell signalling pathways responsible for growth. Growth driving cell signalling, however, is transiently up-regulated regionally during muscle injury and repair to activate quiescent satellite cells responsible for muscle regeneration (Gill et al., 2010). Activation of signalling pathways requires the appropriate expression of ligands as well as their receptors and appropriately sulfated or de-sulfated co-receptors. However, ligands are often sequestered in the extracellular matrix (ECM) which need to be liberated for them to activate their receptor. We have previously shown that de-sulfation of the ECM mediated by SULF1/SULF2 enzymes regulates signalling pathways during development, regeneration and disease (Dhoot, 2012; Dhoot et al., 2001; Gill et al., 2016; Gill et al., 2014).

A number of studies have shown that both the Wnt and FGF signalling pathways become activated in ageing muscle (Bernet JD, 2014; Brack AS et al., 2007; Chakkalakal JV, 2012) which have been postulated to satellite cells losing their self-renewal properties as well as inducing cell fate changes that promote them to differentiate into fibroblasts and thus increase muscle fibrosis. Here we firstly examined the expression of SULF1/SULF2 in aged muscle as these enzymes have been shown to inhibit RTK cell signalling but facilitate Wnt signalling during development and disease (Ai et al., 2003; Lai et al., 2003; Rosen and Lemjabbar-Alaoui, 2010; Sahota and Dhoot, 2009). We show that expression of SULF1/SULF2 increased in muscle with age. Furthermore, this correlates with the onset of spontaneous satellite cell activation.

We also examined the expression of SULF1/SULF2 as well as signalling molecules in a mouse model of accelerated ageing. Ercc1Δ/- hypomorphic mutant mice progressively show signs of ageing in all organs from about 8 weeks of age which is much more severe than in geriatric wild-type mice. Ercc1Δ/- mutant mice die at 4-6 months of age. This model is valuable in that it not only displays most features of normal aging but is amenable to experimentation in a timely manner. Here we show that not only does the muscle of Ercc1Δ/- show elevated levels of SULF1/SULF2 as well as Wnt molecules but that their expression can be significantly decreased by an intervention that promotes muscle survival and growth.

## RESULTS

### SULF1 and SULF2 expression in developing, regenerating and ageing muscle

SULF1 and SULF2 are barely detectable in normal 2-6 month old mouse muscles using immunocytochemical or qPCR procedures (Figure 1) with the exception of some rare isolated activated satellite cells. This follows a high level of SULF1 and SULF2 expression during early fetal muscle growth with their re-activation in adult only during muscle regeneration in response to injury or disease as is observed in mdx mouse muscles undergoing muscle regeneration (Figure 1). Restriction of SULF1/SULF2 enzymes to myogenic cells in such regenerating areas was confirmed by their correlation with staining for skeletal muscle myosin heavy chains (Figure 1.I & J). Spontaneous SULF1 and SULF2 re-activation in adult muscle, however, gradually increases with age as significant levels of these enzymes are detectable at 28 months of age not only in isolated activated satellite cells but also more wide expression on the cell membrane (Figure 1.G, H, K, L) using both immunocytochemical and qPCR procedures. Quantification by qPCR procedure showed relatively higher levels of Sulf1 when compared with Sulf2 (Figure 1.M & N) although presence of both these enzymes was easily apparent by fluorescent antibody staining for SULF1 and SULF2.

**FIGURE 1.**
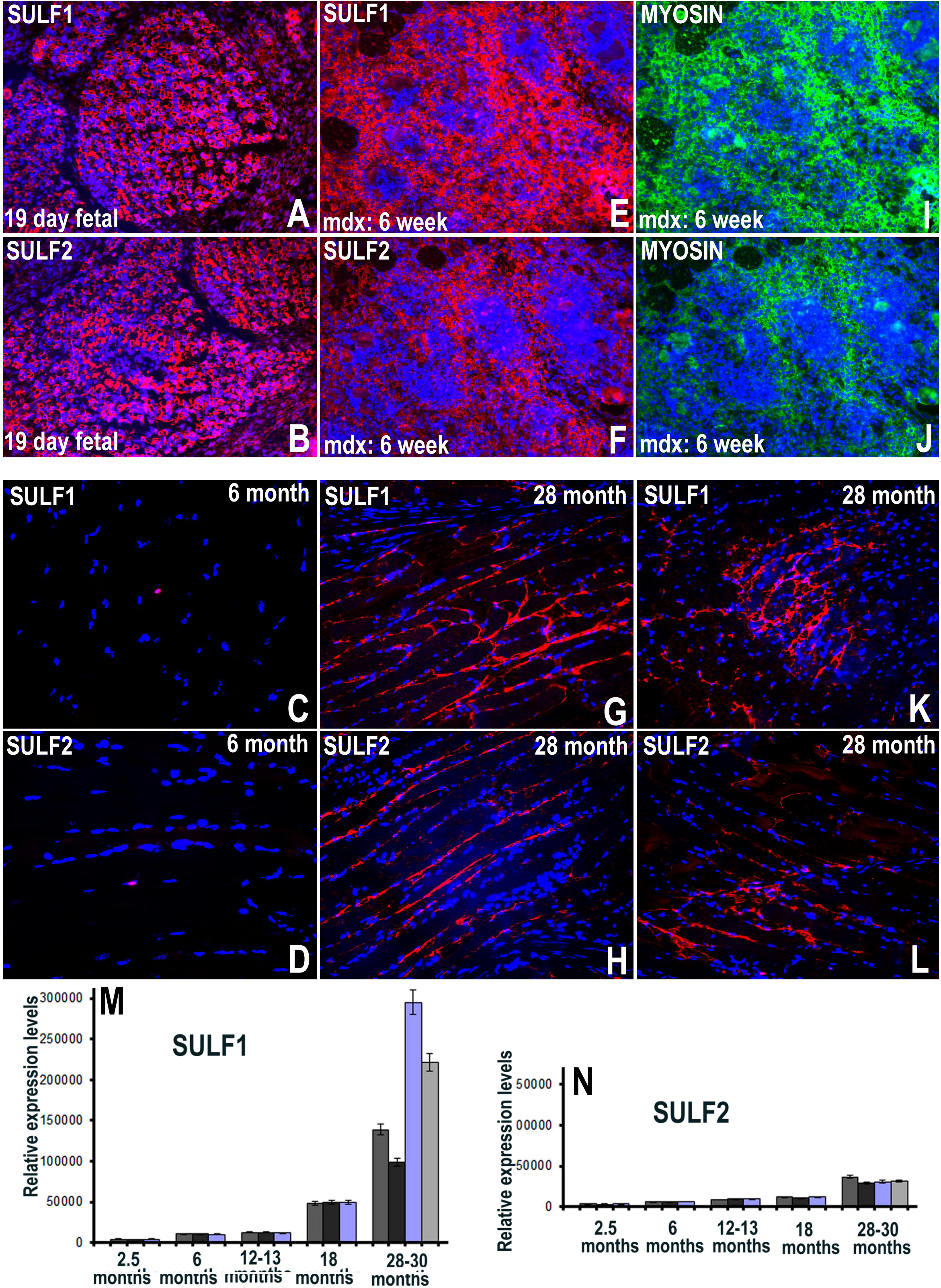
Pattern of SULF1 and SULF2 expression in fetal (A, B) and 6 month (C, D) of Gastrocnemius muscle was compared with their expression pattern in mdx regenerating (E, F) and 28 month old aged Gastrocnemius muscle (G,H, K, L). Sulf1 and Sulf2 expression in 6-week old mdx regenerating muscle (E, F) is also compared with myosin heavy chain expression (I, J) using immunofluorescence staining. Procedure of qPCR was used to measure the changes in the levels of Sulf1 (M) and Sulf2 (N) in this muscle at different ages.

### Re-activation of receptor tyrosine kinase (RTK) and Wnt signalling in ageing muscle

The re-activation of SULF1/SULF2 in ageing muscle prompted us to investigate the re-activation of signalling pathways known to be modulated by SULF enzymes in a positive or a negative manner. As was the case with SULF1 and SULF2, the presence of phospho-cMet and phospho-FGFR1 was considerably increased in 28 month old mice when compared with their barely detectable levels restricted only to an occasional activated satellite cell in 6-month old mice (Figure 2). The levels of these phosphorylated receptors showed intermediate level of increase at 18 months of age (Figure 2. B & E).

**FIGURE 2.**
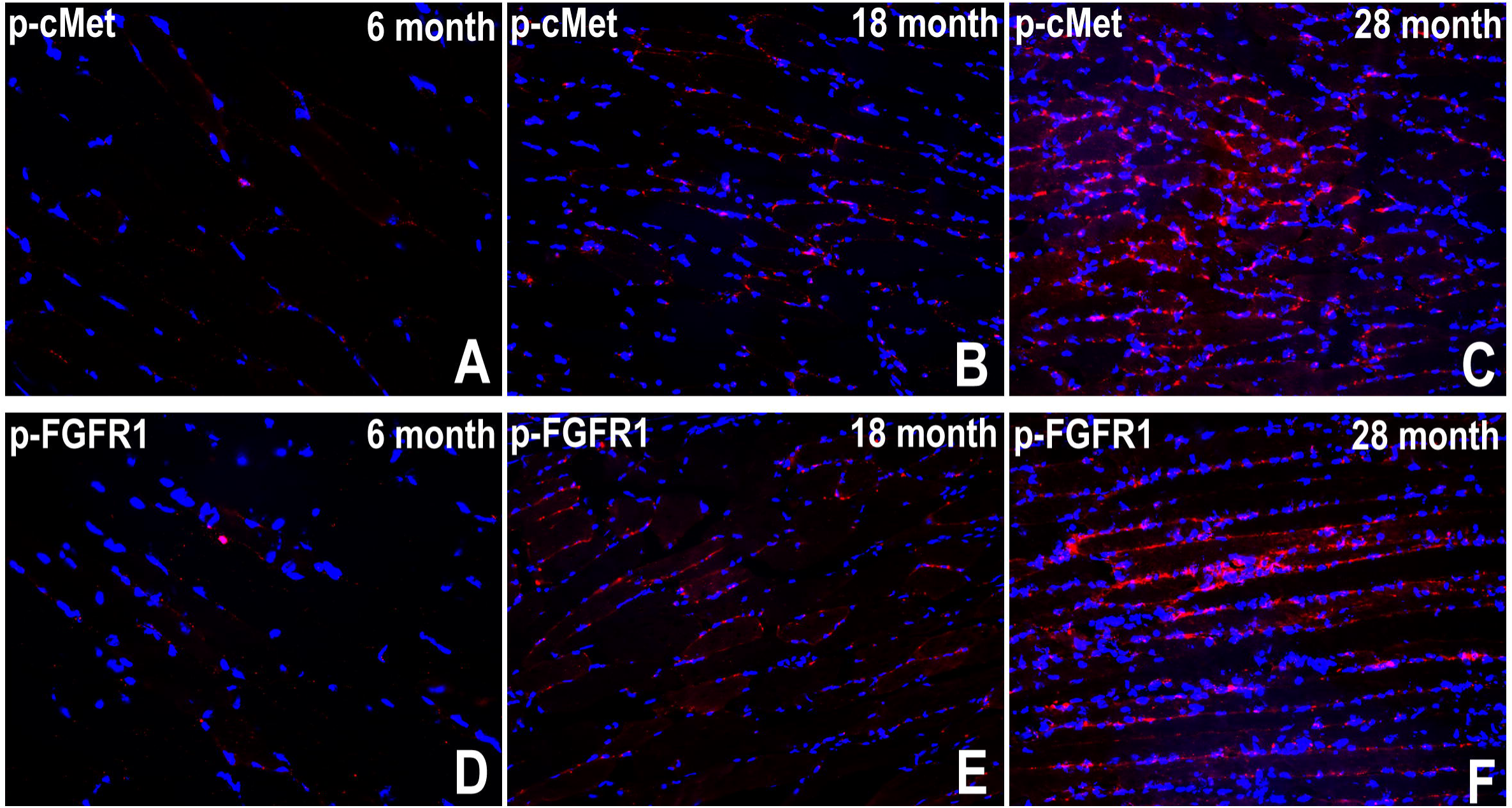
Progressive increase in the levels of phospho-cMet (A,B,C) and phospho-FGFR1 (D,E,F) in Gastrocnemius muscle with age investigated at 6 months (A, D), 18 months (B, E) and 28 months (C, F) using immunocytochemical staining procedure.

In ageing muscle, we also examined the activation of Wnt signalling known to be promoted by SULF1/SULF2 enzymes and well recognised to regulate muscle development and regeneration (Dhoot et al., 2001). Activation of Wnt signalling in younger and ageing muscle was compared using antibodies to Wnt1, Wnt2, Wnt3a, Wnt4, Wnt6 and wnt7a ligands as they can regulate muscle growth (Gill et al., 2010; Hitchins et al., 2013) in a positive and negative manner. Comparison of 28 month old Gastrocnemius muscle with 6 month old muscle shows marked increase in Wnt signalling during later stage (Figure 3). Aged muscle at 28 month stage showed activation of all these Wnt ligands although the level of increase was greatest for Wnt1 and Wnt2 with lower level increases in other Wnts, particularly Wnt7a showing the lowest level of expression (Figure 3). The level of increase in Wnt3a, Wnt4 and Wnt6 in such ageing muscle was relatively moderate (Figure 3).

**FIGURE 3.**
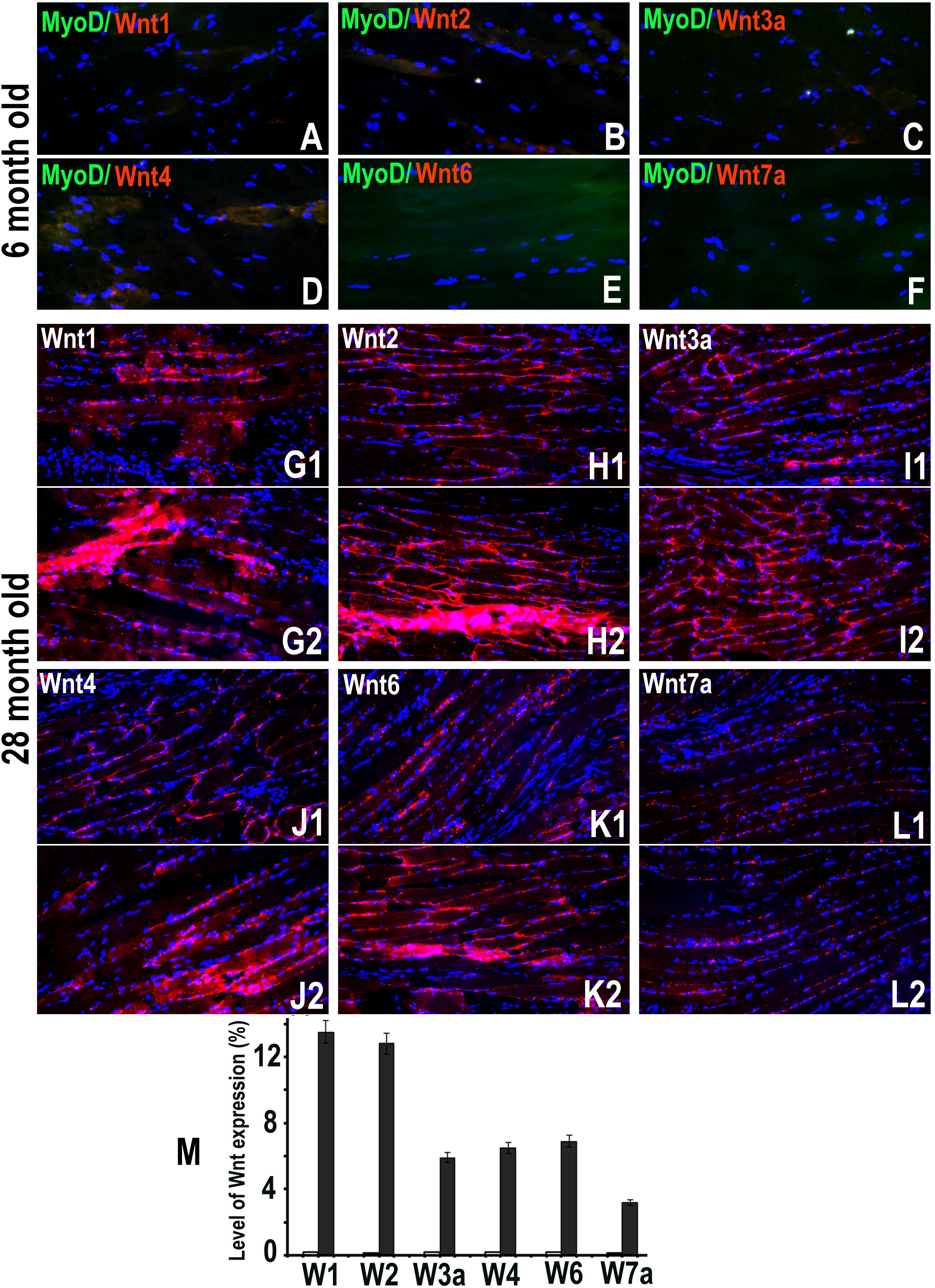
Progressive increase in the levels of several Wnt ligands (Wnt1, Wnt3a, Wnt4, Wnt6, Wnt71) with age in Gastrocnemius muscle is apparent when levels are compared at 6 (A-F) and 28 months (G-L) of age using immunocytochemical staining procedure. Some variation in the levels of Wnts was also observed between individuals of similar age as well as some regional variation (G1/G2, H1/H2, I1/I2, J1/J2, K1/K2, L1/L2) in the same muscle. Quantification of staining levels using volocity software showed higher increase in the levels of Wnt1 and Wnt2 with lowest increase in Wnt7a and moderate levels of increases in Wnt3a, Wnt4 and Wnt6 (M).

### Relative level of satellite cell activation, cell proliferation and response to HGF in ageing muscle *in vitro*

The level of *in vitro* spontaneous satellite cell activation was further examined by preparing single muscle fibres for immediate analysis following isolation i.e time 0hr. Unlike the early developmental stages of 2-3 months showing rare or no activated satellite cells at 0hr, single fibres isolated from later stages showed the presence of a number of activated satellite cells at time 0 identified by their larger nuclear size and SULF1/SULF2 and or MyoD staining. While the number of activated satellite cells at a later stage of 12 months generally showed the activation of some individual satellite cells, single fibres from 2 year old muscles also showed some 2 or 4-cell clusters at 0hr (Figure 4) although not all cells in such clusters showed SULF or MyoD activation.

**FIGURE 4.**
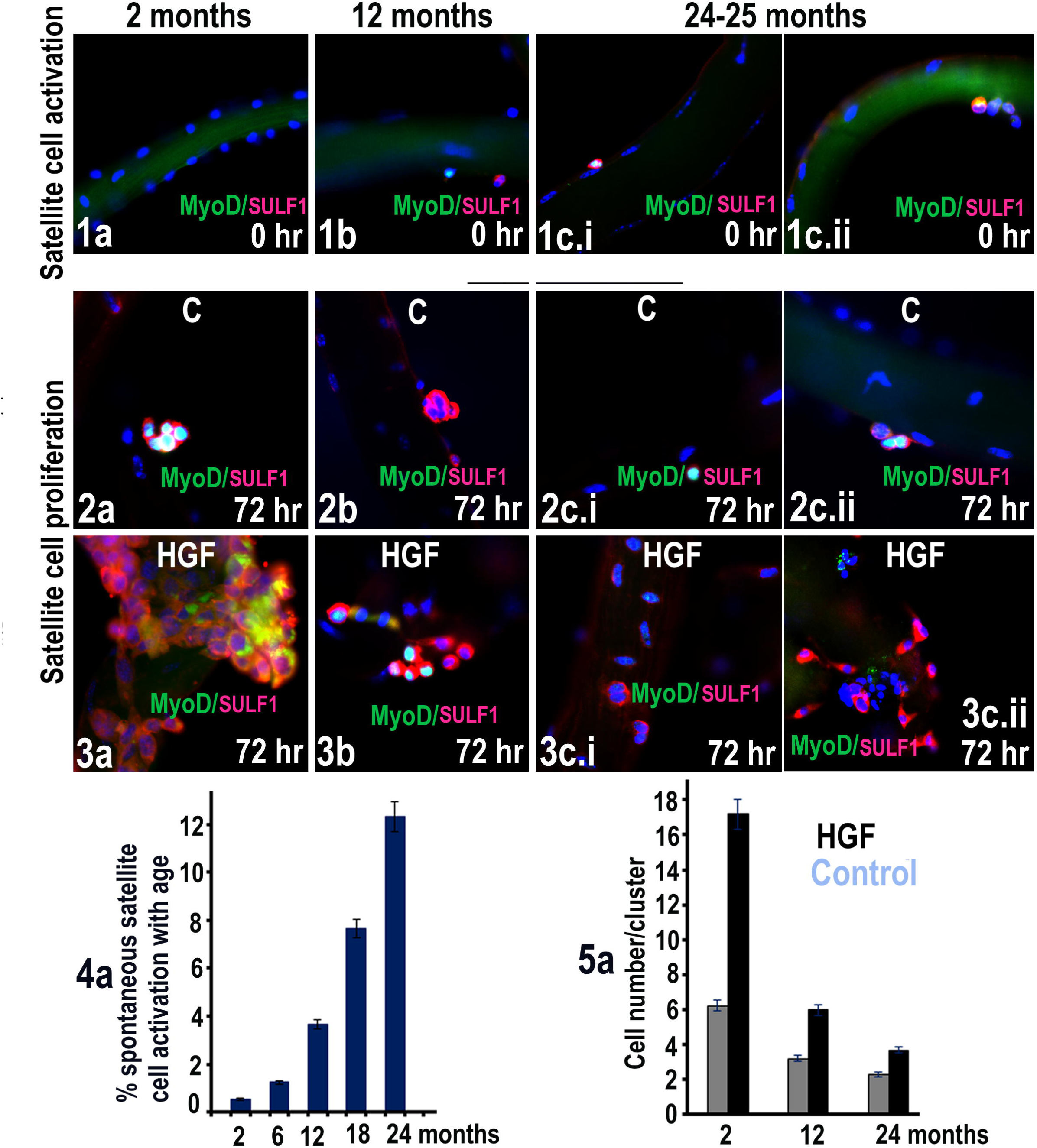
Sulf1 (red) and MyoD (green) immunocytochemical staining of single muscle fibres at 0hr following isolation to determine level of spontaneous satellite cell activation with age (1a,1b,1c.i and 1.c.ii) analysed in EDL muscle at 2, 12 and 24-25 months of age. The proliferative potential of satellite cells on single fibres prepared from 2 month, 12month and 24-25 months was further investigated by in vitro culture of muscle fibres for 72hr without (2a,2b,2c.i and 2c.ii) and with HGF (3a,3b,3c.i and 3c.ii). Relative levels of spontaneous satellite cell activation at 2, 6,12,18 & 24 months and changes in proliferative potential at 2, 12 and 24 months is shown in 4a and 5a.

The *in vitro* culture of isolated single fibres for 72 hours showed reduced rate of satellite cell proliferation apparent from the reduced size of cell clusters on fibres isolated from 2-year old mice in comparison with cell cluster size on fibres from 2.5 month old mice and intermediate size for 12 month old mice (Figure 4).

Addition of HGF to single fibre cultures increases satellite cell proliferation but HGF response of single fibres from older mice in comparison with younger mice showed a much lower level of satellite cell proliferation (Figure 4).

### Earlier re-activation of SULF1/SULF2 and cell signalling in younger Ercc1Δ/- progeric mouse muscles with accelerated ageing but their attenuation in sActRIIB treated muscles

To determine if cell signalling and spontaneous satellite cell activation was enhanced or earlier in progeric mouse muscles with accelerated ageing, we investigated the age-related onset of SULF1/SULF2 and cell signalling in the Gastrocnemius muscle of Ercc1Δ/- mice at 16 weeks of age. Unlike the muscles from 16 week old WT mice that show no overt cell signalling or SULF1/SULF2 activation at this stage, their activation in age-matched Ercc1Δ/- mouse muscles was clearly apparent in regions showing some muscle damage identified from their H&E and immunocytochemical staining for Laminin (Figure 5). This was also apparent from Laminin staining surrounding not only large muscle fibres but also encircling some very small cells that did not appear to grow in size as intermediate size myogenic cells were usually not apparent in such regions (Figure 5G). Such areas also demonstrated activation of cell signalling as for example is revealed by phospho-FGFR1 staining in progeric but not in age-matched WT mouse muscles (Figure 6). Further immunocytochemical staining analysis also demonstrated that SULF1/SULF2 activation was not restricted to only the areas that appeared damaged but also observed on the cell membranes of the apparently uninjured muscle fibres (Figure 6).

**FIGURE 5.**
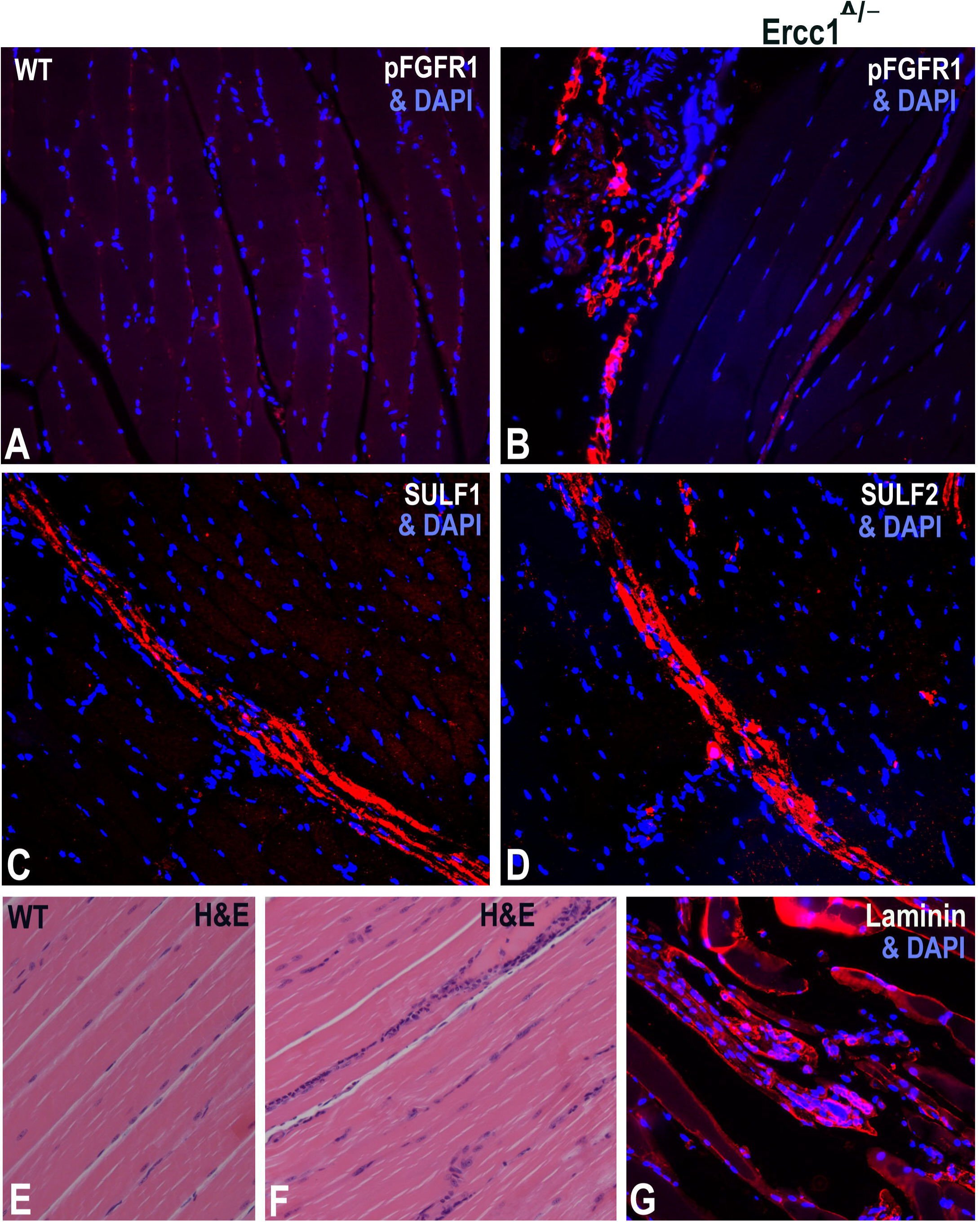
Sulf1/Sulf2 or cell signalling components are not detectable in significant amounts in wild type mouse Gastrocnemius at 16 weeks of Age as shown for pFGFR1 in A. The regional staining for pFGFR1, SULF1 and SULF2, however, is apparent in age matched Ercc1Δ/− muscle. Such areas of positive staining could represent areas of damaged muscle fibres and regenerating myotubes in Ercc1Δ/− muscle as is apparent from H&E (F) and Laminin (G) staining but not from H&E staining of WT muscle (E).

**FIGURE 6.**
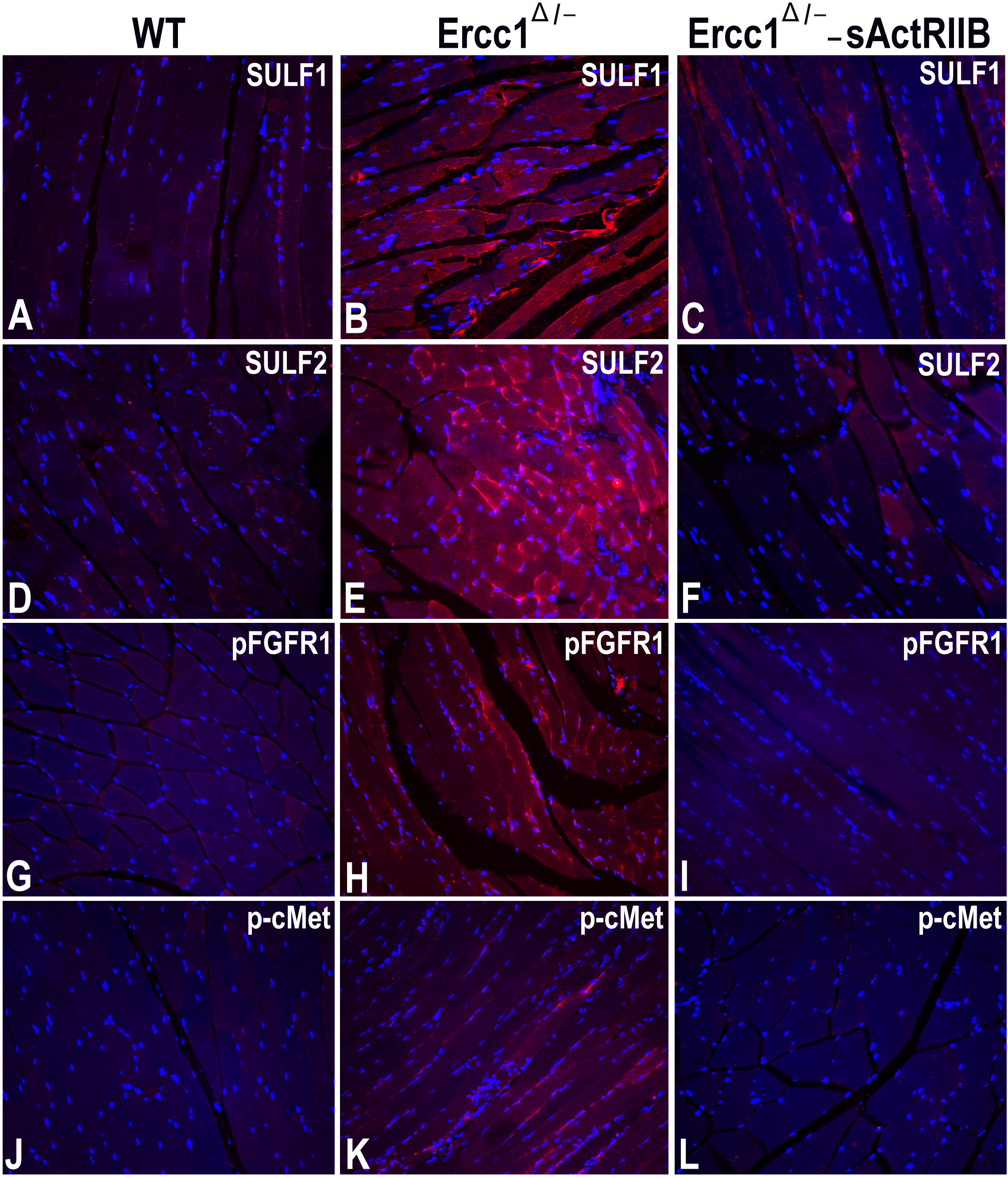
Unlike wild type controls (A, D, G & J), SULF1 (B), SULF2 (E), pFGFR1 (H) & p-cMet (K) activation became detectable in a proportion of 16 week old apparently uninjured Ercc1Δ/− Gastrocnemius muscle fibres. The expression of SULF1 (C), SULF2 (F), pFGFR1 (I) & p-cMet (L), however, was barely detectable in sActRIIB treated Ercc1Δ/− muscles that show activation in only an occasional activated satellite cell.

Previously we have shown that attenuating Myostatin/Activin signalling significantly improved both the quality and quantity of muscle in the Ercc1Δ/- progeric mouse (Alyodawi K et al., 2019). Having shown the aberrant expression of SULF1/SULF2 in the muscle of the progeric mice, we next determined the impact of dampening down Myostatin/Activin signalling on their expression. Remarkably muscles treated with the soluble Activin Receptor Type IIB ligand trap (sActRIIB) for 9 weeks displayed markedly decreased levels of SULF1/SULF2. Only a few activated satellite cells similar to earlier younger muscles showed some SULF1/SULF2 staining but no cell membrane staining was observed in such muscles.

We also investigated if RTK or Wnt signalling was activated in Gastocnemius muscles of 16 week old Ercc1Δ/- mouse muscles since activation of several Wnts was observed in ageing WT muscles that was particularly pronounced during later stages of ageing (Figure 7). Similar to older WT mouse muscles, activation of Wnt signalling in younger Ercc1Δ/- mouse Gastrocnemius was observed at an earlier stage of 16 weeks. As was the case for SULF1/SULf2, cMet and FGF signalling, attenuation of activin signalling by treatment of Ercc1Δ/- progeric mice with activin ligand trap over 9 weeks, also considerably reduced Wnt, cMet and FGF signalling in such treated mice (Figure 7).

**FIGURE 7.**
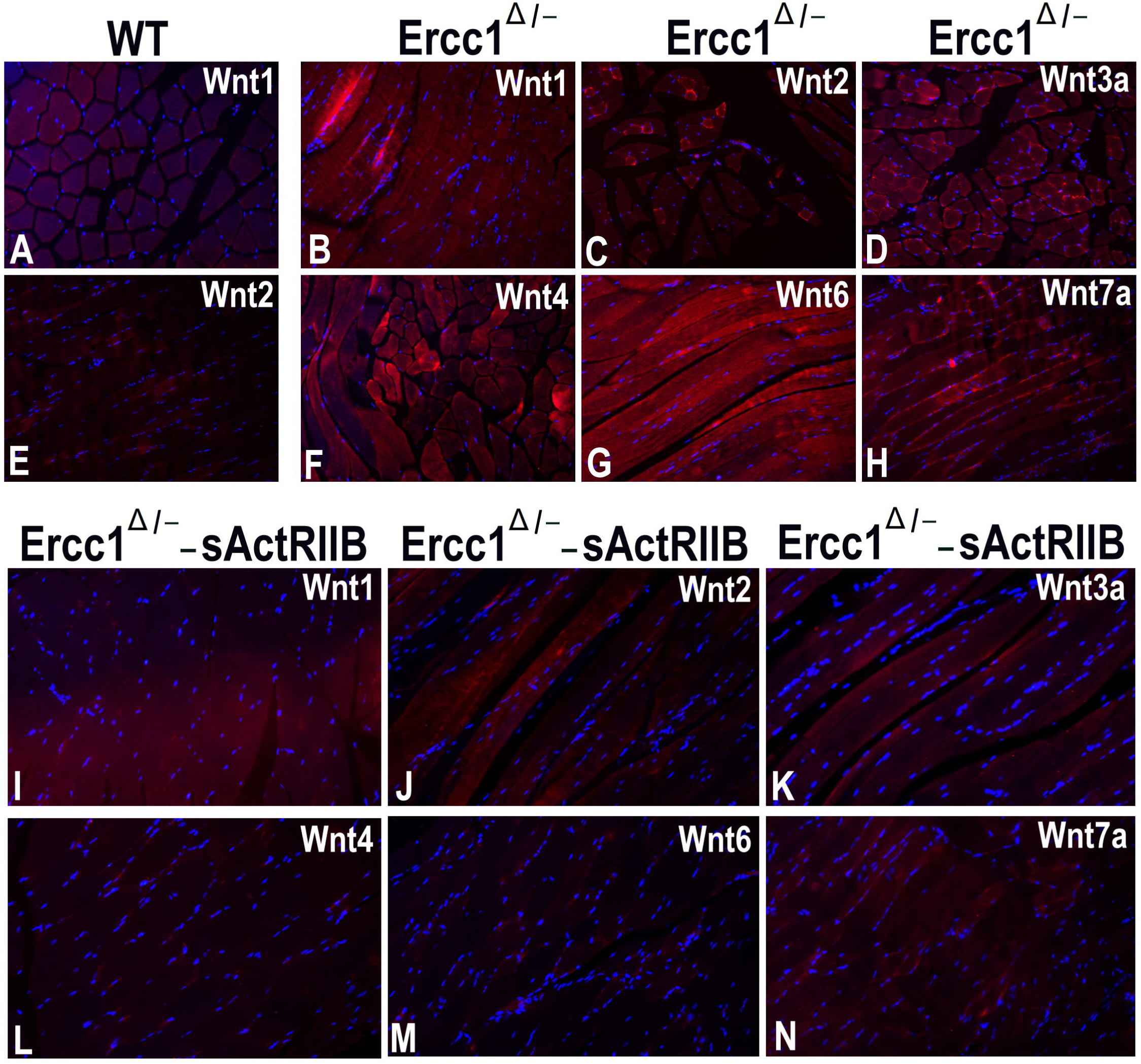
Unlike wild type controls (A & E), Wnt signalling detected by the expression of Wnt ligands Wnt1, Wnt2, Wnt3a, Wnt4, Wnt6 and Wnt7a (B-D, F-H) was apparent in Gastrocnemius muscles of Ercc1Δ/− mice at16 weeks of age. These Wnt ligands, however, were barely detectable in sActRIIB treated Ercc1Δ/− Gastrocnemius muscles (I-N).

## DISCUSSION

Cell signalling is required for muscle development and growth as well as during regeneration in response to injury. These same signalling pathways, however, are thought to be responsible when aberrantly activated during chronic injury and during the ageing process. Here we show that the expression of Sulf molecules is very high during muscle development and in regenerating muscle. However, the expression of these molecules as well as signalling ligands and receptors then decreases dramatically in adult mouse muscle. The low level expression of these molecules then increases gradually with age until they are expressed at high levels again in 28-month old mouse muscle which corresponds to geriatric age in humans (García-Prat L and P., 2017). We additionally show that the Ercc progeric mouse also displays high levels of Sulf as well as RTK & Wnt molecules at 16-weeks of age, a time point when this mouse displays numerous signs of ageing including sarcopenia.

We have previously shown that Sulfs support myogenesis by promoting Wnt signalling; by liberating bound ligands through their enzymatic ability to desulfate the HSPGs in the ECM (Gill et al., 2010; Sahota and Dhoot, 2009). Assuming that the relationship between the Sulf/Wnt and muscle cells is conserved during post-natal life then we can develop a scientific rationale for the results presented in this study. We show sporadic expression of Sulfs and Wnts in mononuclear muscle cells. We suggest that here the cells expressing these molecules are in the process of fusing with the muscle fibre as part of the homeostatic pathway to maintain a healthy tissue. Indeed our understanding of the role played by satellite cells during post-natal life has undergone a tremendous revision underpinned by the development of novel microscopic techniques and the generation of novel mouse genetic models. The work of White et al has shown that the myonuclear content of mouse muscle fibres increases up to 3 weeks in mice (White RB et al., 2010). However others have shown that myonuclei continue to be added at least into adulthood and support not only the muscle fibre size but also their muscle phenotype (specific Myosin Heavy chain expression) (Bachman JF et al., 2018; Pawlikowski B et al., 2015). These studies show that satellite cells are activated in un-injured muscle during phases of myonuclei accretion (up to postnatal day 21) and thereafter to replace nuclei that undergo apoptosis. We suggest that the expression of the Sulfs and the signalling components including Wnt ligands that we detected in adult muscle are in those cells that are in the process of differentiating into fusion competent myoblasts and are responsible for maintaining fibre size and muscle phenotype. Therefore at this stage the expression of Sulfs and Wnts is part of a homeostatic process.

In contrast we suggest that in aged muscle the expression of Sulfs, RTK signalling components and Wnts becomes aberrant that plays a part in the muscle losing its normal functional capacity. We present data that shows expression of Sulfs and these signalling components not only in the satellite cells but also in muscle fibres. A similar expression profile has been reported for members of the FGF signalling cascade (Chakkalakal JV, 2012). Herein Chakkalakal and colleagues suggest that the primary function of the FGF molecules is to support fibre repair that relies on autocrine signalling. However, the undesirable side effect of the autocrine action is that it works in a paracrine manner to deregulate the activity of satellite cells. There is accumulating evidence that aberrant signalling activation in satellite cells that leads to them losing their normal function either by compromising their self-renewal capacity or losing their muscle identity by converting into fibroblasts (Bernet JD, 2014; Brack AS et al., 2007; Chakkalakal JV, 2012). We suggest that the expression of Sulfs (as well as Wnts, FGF & cMet) would fit into this model and that the expression in the fibres is desirable but not in the satellite cells. This hypothesis suggests that both FGF and Wnt signalling are required to attenuate age related changes in muscle fibres. Future work using conditional ablation of genes in muscle fibres versus satellite cells could test such a proposal. Dynamic cell signalling regulates the stem cell pool of many tissues, including satellite cells of skeletal muscle to preserve regenerative capacity throughout life although regenerative potential of many stem cells declines with age. It, however, remains to be determined whether such decline relates to the exhaustion of the satellite cell pool or their limited and finite ability to proliferate that limits or reduces muscle repair during later stages of life.

Stem cells are normally activated during repair or only when required and preserve a stem cell sub-population set aside that does not undergo repeated rounds of cell division. A recent study (Chakkalakal JV, 2012), however, reported spontaneous satellite cell activation in the absence of apparent injury due to a change in satellite cell niche e.g. activation of FGF signalling breaking satellite cell quiescence. Such an age-related change in muscle stem cell niche can trigger unwarranted satellite cell activation leading to reduced longer term regenerative capacity of such stem cells. Satellite stem cell niche, however, constitutes complex microenvironment that does not only include FGF cell signalling but many other cell signalling pathways that can equally promote/substitute, inhibit or regulate muscle cell repair in ageing muscle. We investigated the role of SULF1/SULF2 cell signalling in this study that are known to promote some cell signalling pathways such as Wnt signalling while inhibit many receptor tyrosine kinase mediated cell signalling pathways. Raised levels of SULF expression thus have the potential to promote Wnt signalling but inhibit FGF and/or HGF cell signalling. This can thus disrupt satellite cell quiescence as well as unwarranted or regulated cell signalling. Levels of both SULFs were clearly up-regulated during ageing process that our earlier in vitro study showed to be associated with early phase of satellite cell activation and proliferation but down-regulated during later phase of myogenic differentiation (Gill et al., 2010). Concomitent increase in Wnt signalling indicates increased Wnt signalling but yet not sufficient muscle repair. Concomitent increase in RTK cell signalling pathways likely to be inhibited or down-regulated by SULFs also indicates dysregulated cell signalling. Based on SULF1/SULF2 and MyoD staining, the failure of the activated satellite cells to undergo stepwise cell proliferation and differentiation appeared more regulated in satellite cells of younger muscles than satellite cells of older muscles.

What triggers spontaneous cell signalling and satellite cell activation, however, is not clear unless the threshold of satellite cell activation in the ageing muscle changes with age and lowered with time. Unlike fetal and neonatal growth periods with regulated sequential cell signalling pathways ensuring efficient and ordered cell proliferation and differentiation, simultaneous aberrant activation of multiple cell signalling events may disrupt micro-repair in such regions. Persistent satellite cell activation may also limit the proliferative potential of satellite cells programmed to undergo only a finite number of cell cycles. The limited potential of satellite cells from older muscles was clearly apparent from our *in vitro* analyses as the number of satellite cells/cluster was considerably reduced in the aged muscle. A proportion of such satellite cells also appeared morphologically differentiated or ragged. Unlike satellite cells from younger muscle, satellite cells of ageing muscles also did not respond sufficiently to growth factor exposure such as HGF. The older satellite cells with reduced regenerative potential thus become less responsive to growth enhancing factors with age.

Activated satellite cells during earlier *in vivo* as well as *in vitro* stages generally showed single cell activation as satellite cell clusters are generally not apparent until after 24 hours while up to 4-cell clusters were sometimes observed on older muscle fibres at time 0hr. Such in situ proliferation of activated satellite cells should promote muscle cell repair but yet loss of muscle mass is observed in ageing muscle. This may result from dysregulated cell signalling as indicated by non-activation of simultaneous MyoD and SULFs in some activated satellite cells in aged muscle unlike satellite cells of younger muscles (Gill et al., 2010). The finite ability of satellite cells to undergo only a limited number of cell cycles thus limiting cell proliferation may explain loss of muscle mass but it is unclear how activity or exercise helps maintain muscle mass for longer. It is possible that better cell signalling regulation due to maintenance of higher satellite cell activation threshold in exercised muscle helps preserve a set of satellite cells as a reserve population. Role of muscle activity in satellite cell preservation is also supported by studies of Shefer et al (Shefer G1 et al., 2006) showing relatively higher satellite cell pools of older Soleus, a postural muscle versus EDL muscle.

In this study we show that the Ercc model of accelerated ageing shows elevated expression of Sulf1/Sulf2 as well as Wnt molecules and RTK signalling components at an earlier stage as those displayed by aged muscle. Importantly we show that the expression of all factors associated with ageing are in general normalised by the sActRIIB molecule. Previously we have shown that sActRIIB treatment of the Ercc mouse for an identical period as deployed in this study lead to improved intracellular architecture as well as a normalisation of the ECM which we posited was due to increased autophagy (Alyodawi K et al., 2019). In the same study we showed, using the single fibre model, that satellite cell activity was normalised. The results from this study add an extra layer of understanding of how the introduction of sActRIIB promotes muscle health in the progeric model. We suggest that it improves muscle fibre homeostasis so that the fibres never get to the point where they need to express signalling molecules to maintain its function. As a consequence the satellite cells are not aberrantly activated which helps attenuate the ageing process.

## MATERIALS & METHODS

### Mice and Ethical approval

Normal timed ageing muscles of C57BL/6J mice were obtained from mice supplied by Charles River. The experiments were performed under a project license from the United Kingdom Home Office in agreement with the Animals (Scientific Procedures) Act 1986. The RVC and University of Reading Animal Care and Ethical Review Committee approved all procedures. Animals were humanely sacrificed via Schedule 1 killing

Ercc progeric mice and sActRIIB treatment: Control (Ercc1+/+) and transgenic (Ercc1Δ/-) mice were bred as previously described (Alyodawi K et al., 2019) and maintained in accordance to the Animals (Scientific Procedures) Act 1986 (UK) and approved by the Biological Resource Unit of Reading University or the Dutch Ethical Committee at Erasmus MC. Mice were housed in individual ventilated cages under specific pathogen free conditions (20–22°C, 12–12 hr light–dark cycle) and provided food and water ad libitum. Since the Ercc1Δ/− mice were smaller, food was administered within the cages and water bottles with long nozzles were used from around two weeks of age. Animals were bred and maintained (for the lifespan cohort) on AIN93G synthetic pellets (Research Diet Services B.V.; gross energy content 4.9 kcal/g dry mass, digestible energy 3.97 kcal/g). Post-natal Myostatin/Activin block was induced in seven week-old males, through intraperitoneal injection (IP) with 10 mg/kg of soluble Activin receptor IIB (sActRIIB-Fc) every week two times till week 16 (Alyodawi K et al., 2019). Each experimental group consisted of a minimum of 5 male mice. Lifespan experiments were performed on both genders.

### CELL CULTURE

EDL muscles were carefully dissected from C57BL/6J mice of different ages following Home-Office-approved Schedule 1 killing procedure. Each dissected muscle was transferred to a bijou containing 1 ml of 0.12% Type 1 collagenase (Sigma) in DMEM. Isolated muscles in collagenase solution were incubated at 37°C for 2-3 hours with light shaking every 15 minutes, until the muscle fibres had dissociated. Once dissociated, the collagenase activity was inhibited by the addition of 10% FCS**-**DMEM before single-fibre selection was carried out under a Leica stereo microscope and 12-15 single fibres transferred into each of the wells of a 24-well Linbro plate with 0.5 ml/well culture medium. The normal culture medium was composed of DMEM (Gibco), 4 mM L-glutamine (Sigma), 1% penicillin-streptomycin (Sigma), 10% foetal calf serum (FCS) and 0.5% chick embryo extract (MP Biomedicals). HGF at 100 ng/ml in some experiments as specified was added at time 0. The dissociated single muscle fibres suspended in culture medium were incubated at 37°C in 5% CO2 after 72 hours while a proportion of isolated muscle fibres were fixed in 4% paraformaldehyde for 15 minutes without incubation at this stage of 0hr. Following immunostaining (Gill et al., 2010; Otto A et al., 2008) and imaging, the total number of satellite cells per cluster were counted for each category at 0 and 72 hours. A minimum of 30 cell clusters from at least 15 different muscle fibres were counted for each group. The data from multiple clusters were pooled to obtain a mean (± s.e.m.) for each category. Data are presented as mean ± standard deviation. Statistical analysis was performed using ANOVA, regarding *P*<0.05 as statistically significant.

### Immunocytochemistry

Single muscle fibres following 15 minute fixation in 4% paraformaldehyde and PBS washes were incubated with permeabalisation buffer for 15 min at room temperature before PBS washes and incubation with 10% FCS to block non-specific antibody binding as previously described (Gill et al 2010). Fibres were then incubated with primary antibodies against MyoD (1/100) and SULF1 (1/200) or SULF2 (1/100). Goat anti-rabbit immunoglobulins secondary antibodies and both fluorochromes were diluted 1/400. All primary antibody reactions were incubated overnight at 4 °C followed by secondary antibody incubations for 1 hour each at room temperature. The binding of rabbit primary antibodies was detected using streptavidin Alexa Fluor 594 or Alexa Fluor 488 fluorochrome bound to biotin-linked goat anti-rabbit immunoglobulins. The binding of MyoD mouse immunoglobulins was detected using goat anti mouse immunoglobulins linked to 488 fluorochrome. Following four PBS washes between and after each incubation, labelled tissues were mounted in polyvinyl alcohol mounting medium with DABCO and 2.5 μg/ml DAPI for nuclear visualisation and photographed using a Leica DM4000B fluorescent microscope. Paraffin tissue sections were also stained using single or double immunofluorescence procedure. Tissue sections treated with permeabalisation buffer for 15 min at room temperature, PBS washes and incubation with 10% FCS were treated with diluted primary and secondary antibodies as described above. Sections treated with pre-immune rabbit sera (not shown) were similarly incubated with fluorochrome-labelled secondary antibodies as controls. The dilutions of other primary antibodies were as follows: rabbit anti Laminin: 1/50, rabbit anti phospho-FGFR1: 1/100, rabbit anti phospho-cMet: 1/100, rabbit anti Wnt1: 1/100, rabbit anti Wnt2: 1/100, rabbit anti Wnt3a: 1/300, rabbit anti Wnt4: 1/100, rabbit anti Wnt6: 1/100, rabbit anti Wnt7a: 1/100 and mouse (83B6) anti myosin heavy chain: 1/100.

### qPCR

RNA from mouse Gastrocnemius muscles of different ages was prepared using an Invitrogen Trizol method. RNA, 1μg from each sample was reverse-transcribed with SuperScript II RNase reverse transcriptase (Invitrogen) using random primers. Real-time qPCR was carried out as previously described (Zaman et al., 2016) using QuantiTect SYBR green PCR kit and Opticon 2 LightCycler (MJ Research, Waltham, MA, USA). The expression levels of Sulf1 and Sulf2 were normalised to the reference gene 18s rRNA. PCR primers used in this study were as follows: Sulf1 (5′-ATGAAGTATTCCCTCTGGGCTCTG-3′; 5′-CAATGTGGTAGCCGTGGTCC-3′); Sulf2(5′-ATGGCACCCCCTGGCCTGCCACTAT-3′; 5′-CATAGACTTGCCCTTCACCAGCCC-3′) and 18s RNA (5′-CGCGGTTCTATTTTGTTGGT-3′; 5′-AGTCGGCATCGTTTATGGTC-3′).

